# PTMscape: an open source tool to predict generic post-translational modifications and map hotspots of modification crosstalk

**DOI:** 10.1101/257386

**Authors:** Ginny X.H. Li, Christine Vogel, Hyungwon Choi

## Abstract

While tandem mass spectrometry can now detect post-translational modifications (PTM) at the proteome scale, reported modification sites are often incomplete and include false positives. Computational approaches can complement these datasets by additional predictions, but most available tools are tailored for single modifications and each tool uses different features for prediction. We developed an R package called PTMscape which predicts modifications sites across the proteome based on a unified and comprehensive set of descriptors of the physico-chemical microenvironment of modified sites, with additional downstream analysis modules to test enrichment of individual or pairs of modifications in functional protein regions. PTMscape is generic in the ability to process any major modifications, such as phosphorylation and ubiquitination, while achieving the sensitivity and specificity comparable to single-PTM methods and outperforming other multi-PTM tools. Maintaining generalizability of the framework, we expanded proteome-wide coverage of five major modifications affecting different residues by prediction and performed combinatorial analysis for spatial co-occurrence of pairs of those modifications. This analysis revealed potential modification hotspots and crosstalk among multiple PTMs in key protein domains such as histone, protein kinase, and RNA recognition motifs, spanning various biological processes such as RNA processing, DNA damage response, signal transduction, and regulation of cell cycle. These results provide a proteome-scale analysis of crosstalk among major PTMs and can be easily extended to other modifications.

**Contact:** all correspondence should be addressed to.

## Introduction

Protein post-translational modifications (PTMs) regulate cellular functions in various ways: catalyzing enzymatic activities, conferring substrate specificity to control allosteric interactions, mediating interactions with other molecules such as DNA, co-factors, and lipids, and localizing proteins to organelles^1^. With advances in enrichment techniques for PTMs, high-resolution mass spectrometry (MS) has now become the method of choice to experimentally detect and quantify major modifications at a proteome scale^2^. A wealth of PTM data arising from tandem MS/MS experiments has been curated and shared in public databases such as PhosphoSitePlus (PSP)^3^, PHOSIDA^4^, and Uniprot^5^, and some major modifications such as phosphorylation and ubiquitination have been mapped for multiple species. For instance, as of December 2017, the PSP database described ~240,000 phosphorylation and ~22,000 ubiquitination sites for >20,000 different human proteins.

A comprehensive map of diverse modifications can help us infer not only the role of individual PTMs, but also the complex code of different modifications localized to the same protein jointly modulating biochemical functions through positive and negative regulatory interactions, also known as the PTM crosstalk^6^. A well-known example for such crosstalk is the tumor suppressor gene p53 whose abundant and diverse modifications affect the protein’s activity and subsequent cancer formation.^7–9^ p53 has at least 20 known phosphorylation sites and multiple modifications including acetylation, ubiquitination, methylation, and O-GlcNAc sites. Therefore, a critical first step is to map modification sites across the entire proteome in an unbiased manner, attaching confidence scores that allow removal of false-positive identifications and incorrect site localizations.

However, even for major PTMs, including acetylation, methylation, and glycosylation, experiments with new enrichment techniques^10^ often identify novel modification sites, implying that we have not yet reached the coverage to provide the total modification landscape. While experimental efforts are slowly completing this landscape, there is a growing need for statistical frameworks that integrate these diverse modifications at proteome-scale into a unified, comprehensive PTM map with a minimal number of false positives. In addition, such a map should annotate each modification type by predictive descriptors such as physicochemical properties, protein structure information, and sequence motif information such as position specific amino acid propensity (PSAAP).

In the literature, there already exists a plethora of computational prediction methods for major modifications, where the majority of methods relies on complex machine learning algorithms in combination with sequence-level scoring functions that test for single types of modifications.^11^ The prediction methods vary by the type of algorithms, the use of adjacent residues around candidate sites, the use of three dimensional structure information, and specificity with respect to kinase families (in the case of phosphorylation). Recently, a number of prediction tools has also been reported for ubiquitination^12–14^ or arginine methylation^15^, in which predictions are made based on amino acid properties rather than sequence characteristics due to the lack of clear motifs.

Although these tools provide small-scale predictions in a user-friendly web interface, only a few tools provide batch prediction capability for the whole proteome due to high computational burden by non-linear prediction algorithms – and are therefore incapable of extracting examples like that of p53 in a comprehensive, genome-wide manner. Moreover, each single PTM prediction tool uses a different set of descriptors (predictive features) and the models are trained using different training PTM data, often leaving the user no freedom to provide input to the construction of the prediction models. More importantly, as these methods rely on non-linear prediction algorithms such as SVM with the radial kernel, it remains elusive how each descriptor or a combination of descriptors contributes to the probability of PTM events, challenging the interpretation of the best predictors. Other omnibus tools such as ModPred^16^ perform batch predictions for multiple modifications, but their prediction has limited sensitivity in the whole proteome scale (see below). In addition, the tools’ predictions are made from a pre-trained model which the user cannot modify or rebuild. Further, whole proteome-scale predictions are very time consuming in a standard computing environment, precluding the execution of such analysis for lay users.

To address these limitations and enable researchers to map a variety of modification types across the proteome, here we present PTMscape, a unified, highly sensitive and specific framework for high confidence PTM predictions in whole proteome scale. PTMscape has several key advances. First, it is generically applicable to any PTMs and enables the user to train and test predictions using a comprehensive set of descriptors, while operating both at the whole proteome scale and in a time-efficient manner. Unlike most existing tools, PTMscape provides a full set of precompiled features and facilitates the construction of new training and test data, allowing the user to control the model building process. With these resources, PTMscape achieves prediction accuracy comparable to the best single-modification tools currently available. Second, PTMscape performs further downstream statistical enrichment analysis of individual modifications and their crosstalk within protein domains and within function categories. Third, PTMscape offers model training and prediction for large-scale prediction via local installation of the software. PTMscape is packaged for the popular R environment (http://cran.r-project.org) and the user can take full control to create appropriate training and test data for an entire proteome.

To our knowledge, PTMscape is the first open-source, comprehensive statistical framework of this kind. To demonstrate the utility of this tool, we applied PTMscape to ~17,000 human protein sequences and predicted ~39,000 additional post-translational modifications of five different types, revealing the potential hotspots of extensive crosstalk among the modifications.

## Results

### PTMscape uses linear SVMs and a comprehensive set of predictors to evaluate five major modification types

In PTMscape, we advocate the SVM with linear kernel to perform linear classification to make prediction for each candidate site into a modified site or an unmodified one, while the prediction algorithm can be easily replaced by other methods such as SVM with non-linear kernels or neural networks. In contrast to PTMscape, most prediction tools use SVM with a radial kernel, a nonlinear classification method for flexible and sensitive classification. The rationale for our choice of a linear kernel lies in the ability to interpret the resulting weight coefficients. Although the nonlinear classifier is the more sensitive than the linear counterpart, the decision boundaries of the SVM with a non-linear kernel are too complex for easy interpretation. In addition, a non-linear classification method can be easily over-fitted. In contrast, the linear SVM has the advantage that the optimized weight coefficients directly inform on the contribution of each descriptor to the likelihood of the PTM status.

To evaluate the performance of overall predictions, we first assembled and evaluated a comprehensive set of descriptors which we derived from literature survey. In this initial analysis, we aimed to learn how well the positive sites, i.e. modified sites detected in experiments collected in the PSP database, can be differentiated from the negative sites, i.e. sites not reported in the PTM database, and what information (descriptor) is useful to predict each PTM type.

We compiled descriptors of physicochemical properties in the microenvironment of modification sites from three different sources. The first source consisted of 538 amino acid indexes, which have been previously used in prediction methods, e.g. for ubiquitination^13,17^. The indexes consisted of physicochemical properties for each amino acid,^18^ including, for example, hydrophobicity, propensity to be in secondary structures, free energy change, and residue volume. A large number of these properties were highly correlated with one another. Therefore, we grouped the normalized properties into 53 clusters and used their average values as a representative value for each cluster (see Methods). The second source included residue-specific properties such as access to surface area, half sphere exposure, and the probability of being positioned in secondary structures such as coils, sheets, and helix computed by SPIDER3 software^19^. This information was obtained based on the secondary structure assignment to the protein sequence. The last source was the position specific amino acid propensity (PSAAP) matrix computed for each modification based on experimentally acquired sites (deemed positives).^11^ The final 173 predictors comprised six categories including the average amino acid indexes, four secondary structure features, and the PSAAP scores.

Using PTMscape, we computed these properties for all proteins in the human proteome, for window sizes of 11, 15, and 25 amino acids where the center position contained the modified residue. SPIDER3 was able to assign individual residues to secondary structures for approximately 17,000 proteins, and we perform all our proteome-scale predictions within this set. Following the convention in other publications,^12, 17, 20–22^ we reduced sequence redundancy by removing highly similar sequences, resulting in a total of ~10,700 human sequences considered (see **Methods**). We focused on five different modifications affecting five different residues: phosphorylation (S, T, Y), ubiquitination (K), SUMOylation (K), acetylation (K), and methylation (K, R). We evaluated the performance of PTMscape’s linear SVM classifier for each of the different modification types using 10-fold cross-validation. In the cross-validation, a model was trained on nine folds of the data and tested on the remaining, randomly chosen one-fold, and the same was iteratively applied to all ten folds.

**Table 1** shows the overall prediction performance of linear SVMs across the five different modifications. The area under the curve (AUC) for the receiver-operating characteristic (ROC) was the highest for arginine methylation and lysine SUMOylation (0.79), whereas it was the lowest for lysine ubiquitination (0.64) and acetylation (0.66), for all window sizes (**Supplementary Figure 1**). As expected, the modifications known to have clear global sequence motifs were better predicted than those without a sequence motif, and we discuss this point below. For most modifications, the prediction models using the wide 25 amino acid window performed the best (**Supplementary Table 1**), albeit by a small margin. Hence, we used the 25 amino acid windows hereafter.

**Table 1.** Performance evaluation of the linear SVMs across five modification types. The number of true positive sites used in the 10-fold cross-validation is about half the amount of data present in the PSP database after removal of redundant protein sequences and those that do not have secondary structure information from SPIDER3. AUC - area-under-the-curve; MCC – the highest Matthew’s correlation coefficient at all score thresholds; Sensitivity/Specificity at score threshold corresponding to the highest MCC value.

| PTM type | No. of proteins | No. of PSP sites | Window size 25 |  |  |  |
| --- | --- | --- | --- | --- | --- | --- |
|  |  |  | AUC | MCC | Sensitivity | Specificity |
| acetylation (K) | 3,729 | 10,479 | 0.66 | 0.25 | 0.61 | 0.64 |
| methylation (K) | 1,521 | 2,566 | 0.74 | 0.39 | 0.61 | 0.76 |
| ubiquitination (K) | 4,874 | 22,592 | 0.64 | 0.22 | 0.67 | 0.54 |
| SUMOylation (K) | 1,020 | 2,996 | 0.77 | 0.42 | 0.63 | 0.79 |
| methylation (R) | 2,301 | 5,450 | 0.79 | 0.47 | 0.62 | 0.84 |
| phosphorylation (S) | 8,510 | 76,008 | 0.74 | 0.36 | 0.70 | 0.66 |
| phosphorylation (T) | 6,982 | 28,359 | 0.72 | 0.33 | 0.66 | 0.66 |
| phosphorylation (Y) | 6,097 | 18,645 | 0.70 | 0.30 | 0.72 | 0.58 |

Here the AUC was computed treating all candidate sites not reported in the PSP database as negatives, which should have been the sites which are not modified rather than undetected sites. The assumption, i.e. no detection equals no modification, is currently the only option for construction of a negative set and objective evaluation of proteome-scale predictions. For the reasons discussed above, it is likely that many of these negative sites will be unveiled as positives by future experiments. Therefore, the sensitivity, i.e. the prediction of true positives, calculated here is likely underestimated and better than what is reported in **Table 1**.

The importance of specificity cannot be overstated in view of the prediction capability across different methods in major PTMs. To investigate the root cause of lagging AUC values across most prediction methods, we studied the decision boundaries separating positives and negatives in the feature space more carefully in the context of whole proteome-scale prediction. **Figure 1** shows the plots of Partial Least Squares - Discriminant Analysis, a useful tool for projecting high-dimensional data in a supervised way to separate the groups, across all five modifications using all the descriptors we collected. The figure shows that, with the exception of lysine/arginine methylation and SUMOylation, it is difficult to detect a large number of additional true sites at high specificity based on these features. It is likely due to this challenge that many existing methods resorted to complex non-linear prediction methods, attempting to find the subspace in the feature set that confers increased likelihood of a given PTM event. Nevertheless, the poor overall separation of positives and negatives suggests that it is of paramount importance to maintain a stringent level of specificity in predictions, even at the expense of the sizeable loss in sensitivity. This explains why we choose a score threshold associated with very high specificity rule (99%) in our predictions below.

**Figure 1.**
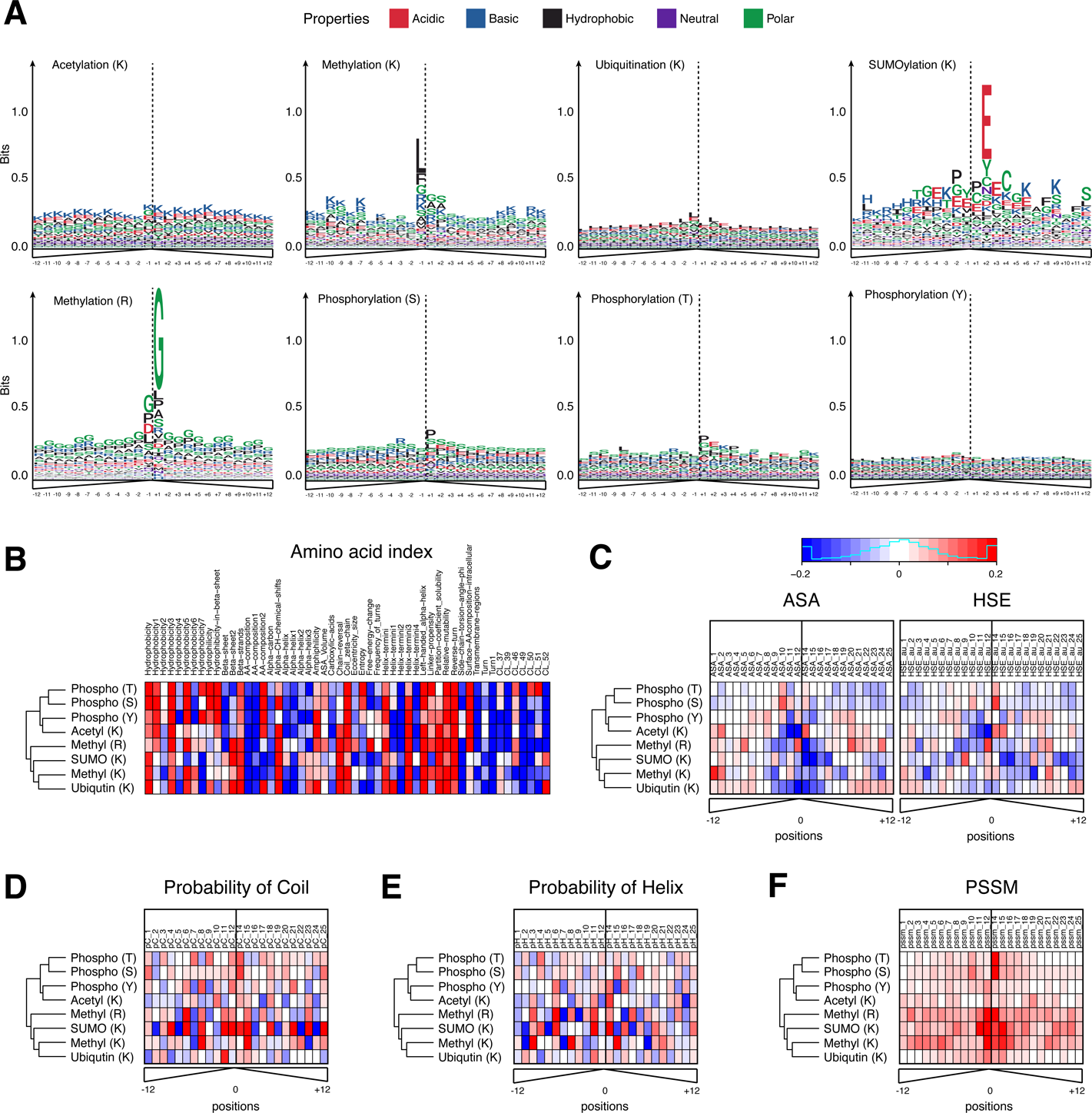
Partial least squares - discriminant analysis plot for five modification types. Red and blue colors indicate positive, experimentally determined modifications sites (those recorded in the PSP database) and negative sites (the remainder of possible sites). The negatives have been randomly sampled to match the same number of positives in each modification to prevent a large number of negatives from masking the positives in each plot.

### Linear SVM is comparable to the best prediction algorithms of individual modifications

Next, we benchmarked the prediction performance of PTMscape’s linear SVM against the algorithms leading in prediction of phosphorylation and ubiquitination, the two modifications most widely studied by mass spectrometry-based proteomics. For phosphorylation, PTMscape’s linear SVM compared very well with the latest kinase-independent phosphorylation prediction tool called PhosphoSVM, which outperforms the majority of global phosphorylation prediction tools.^22^ Using PhosphoSVM’s test data consisting of 9,688 serine, 2,919 threonine, and 1,269 tyrosine positive sites on 2,545, 1,499, and 805 protein sequences, we trained PTMscape’s linear SVMs using predictive variables computed for 25 amino acid windows with 10-fold cross-validation. This setup was identical to that used by PhosphoSVM. Therefore, the only difference between the two methods was that our classifier (predictor) used more features for prediction and a simpler kernel than that of PhosphoSVM. Notably, PTMscape and PhosphoSVM showed similar performance with the exception of serine phosphorylation, where the AUC was greater by 0.03 in PhosphoSVM (**Table 2**).

**Table 2.** Comparison of performance metrics between PTMscape with linear SVM and the best existing prediction methods in phosphorylation and ubiqutination using training and test data sets provided by the latter methods. For PTMscape, we chose score thresholds in three different ways, using the thresholds that give the best MCC, the best F-measure and the same specificity as reported in the benchmarking tool, respectively. The performance metric for other ubiquitination site prediction is from J.-R.Wang *et al*^17^. AUC - area-under-the-curve; MCC - Matthew’s correlation coefficient.

| PTM Type | Methods | AUC | sd AUC | Choice of threshold | MCC | Sensitivity | Specificity |
| --- | --- | --- | --- | --- | --- | --- | --- |
| Phosphorylation (S) | PhosphoSVM | 0.84 | 0.01 | F-measure | 0.30 | 0.44 | 0.94 |
|  | PTMscape | 0.81 | 0.01 | MCC | 0.49 | 0.74 | 0.75 |
|  |  |  |  | F-measure | 0.47 | 0.86 | 0.60 |
|  |  |  |  | Matching specificity | 0.37 | 0.36 | 0.94 |
| Phosphorylation (T) | PhosphoSVM | 0.82 | 0.01 | F-measure | 0.25 | 0.37 | 0.95 |
|  | PTMscape | 0.80 | 0.01 | MCC | 0.48 | 0.71 | 0.75 |
|  |  |  |  | F-measure | 0.46 | 0.85 | 0.60 |
|  |  |  |  | Matching specificity | 0.37 | 0.34 | 0.95 |
| Phosphorylation (Y) | PhosphoSVM | 0.74 | 0.02 | F-measure | 0.21 | 0.42 | 0.87 |
|  | PTMscape | 0.74 | 0.03 | MCC | 0.41 | 0.66 | 0.73 |
|  |  |  |  | F-measure | 0.35 | 0.89 | 0.42 |
|  |  |  |  | Matching specificity | 0.30 | 0.40 | 0.87 |
| Ubiquitination (K) | ESA-ubiSite | 0.95 | n.a | MCC | 0.48 | 0.66 | 0.94 |
|  | ESA-SVM-PCPs | 0.83 | n.a | MCC | 0.27 | 0.76 | 0.75 |
|  | ESA-5NN | 0.65 | n.a | MCC | 0.17 | 0.69 | 0.66 |
|  | Random-SVM | 0.73 | n.a | MCC | 0.17 | 0.67 | 0.67 |
|  | PTMscape | 0.82 | n.a | MCC | 0.29 | 0.64 | 0.83 |
|  |  |  |  | F-measure | 0.28 | 0.43 | 0.92 |
|  |  |  |  | Matching specificity | 0.23 | 0.31 | 0.94 |

PTMscape’s linear SVM classifier also compared very well with the most state-of-the-art ubiquitination prediction tool called ESA-UbiSite.^17^ ESA-UbiSite uses an evolutionary screening algorithm coupled to SVM with non-linear kernel to address the lack of true negatives by iteratively updating the modification status of negative sites in the model-training phase. Using the training data and the high-confidence test data provided by ESA-UbiSite, PTMscape achieved an AUC of 0.82, comparable to the second best algorithm ESA-SVM-PCPs (AUC 0.83) but worse than the best method ESA-UbiSite (AUC 0.95). It is possible that this difference in AUC most likely arose from the specificity calculation, as the test data is very small with only 645 positive sites on 379 proteins. Therefore, although the evolutionary screening algorithm for identifying better negative sites makes valuable contribution, the calculation of AUC remains to be evaluated on a larger test set.

Next, we show that PTMscape outperforms ModPred, which is one of the few available generic tools for modification prediction.^16^ Unfortunately, it was practically infeasible to directly compare PTMscape and ModPred using the same training and test data because we are unable to build the new prediction models for cross-validation within their framework. Therefore, we ran ModPred provided by the developers to predict phosphorylation, ubiquitination, SUMOylation, acetylation, and methylation on ~12,000 nun-redundant protein sequences used in our 10-fold cross-validation scheme, and compared their performance with that of PTMscape. This implies that we made predictions on some of the proteins which were used for training data in ModPred, giving the method a potential advantage over PTMscape. ModPred performed the best with serine phosphorylation at an AUC of 0.74 (**Supplementary Table 2**) followed by threonine/tyrosine phosphorylation, arginine methylation and lysine acetylation with AUC ranging from 0.65 to 0.7. ModPred performed poorly for other lysine modifications (AUC<=0.6). In contrast, the AUCs of PTMscape were better, ranging from 0.64 to 0.74 across all modifications (**Supplementary Table 2**).

### PTMscape’s predictions associated with protein domains and functions

Next, we used PTMscape’s comprehensive descriptor set and linear SVM classification to map five major modifications at proteome scale in a unified statistical framework. We used two-fold cross-prediction (see **Methods**) so that the training data and the test data are completely independent. Predictions were made on the total human proteome, comprising ~17,000 sequences. The predictions were made at score thresholds associated with 99% specificity, in contrast to the conventional choice giving the best Matthew’s correlation or F-measure, to ensure a low false positive rate account for the varying range of AUCs.

**Table 3** shows the experimentally detected sites available in the PSP database (254,116 PTM sites) and the sites newly predicted by PTMscape, which amount to a total of 38,857 new sites. As expected, the number of newly predicted sites compared to the experimentally detected sites varied across the modifications. For modifications with proteome-wide experimental coverage, e.g. phosphorylation and ubiquitination, newly predicted sites accounted for <14% of the sites known from PSP. By contrast, for modifications with incomplete proteome-wide coverage, e.g. SUMOylation and methylation, PTMscape predicted 85~139% new sites even at 99% specificity. Notably, while lysine SUMOylation^23–25^ and lysine/arginine methylation had the most consistent global sequence motifs (**Figure 2A**), they also had many sites with predictive properties that have been unreported in the literature so far, suggesting that our knowledge of these modifications is still incomplete.

**Table 3.** Prediction of five different modifications across the whole human proteome. The two-fold cross-prediction scheme ensured that the prediction model used on a protein sequence did not include any information from the same protein. The ‘ratio’ contains the number of newly predicted sites divided by the number of known sites in the PSP database.

| Modification | Number of modifiable sites | Number of sites in PSP | Number of newly predicted sites | Ratio (Predicted / PSP records) |
| --- | --- | --- | --- | --- |
| acetylation (K) | 515,336 | 17,084 | 5,148 | 0.30 |
| methylation (K) | 528,563 | 3,857 | 5,238 | 1.36 |
| ubiquitination (K) | 497,377 | 35,043 | 4,997 | 0.14 |
| SUMOylation (K) | 526,278 | 6,142 | 5,330 | 0.87 |
| methylation (R) | 511,803 | 7,957 | 5,072 | 0.64 |
| phosphorylation (S) | 650,342 | 108,287 | 6,578 | 0.06 |
| phosphorylation (T) | 436,467 | 45,302 | 4,359 | 0.10 |
| phosphorylation (Y) | 214,390 | 30,444 | 2,135 | 0.07 |

**Figure 2.**
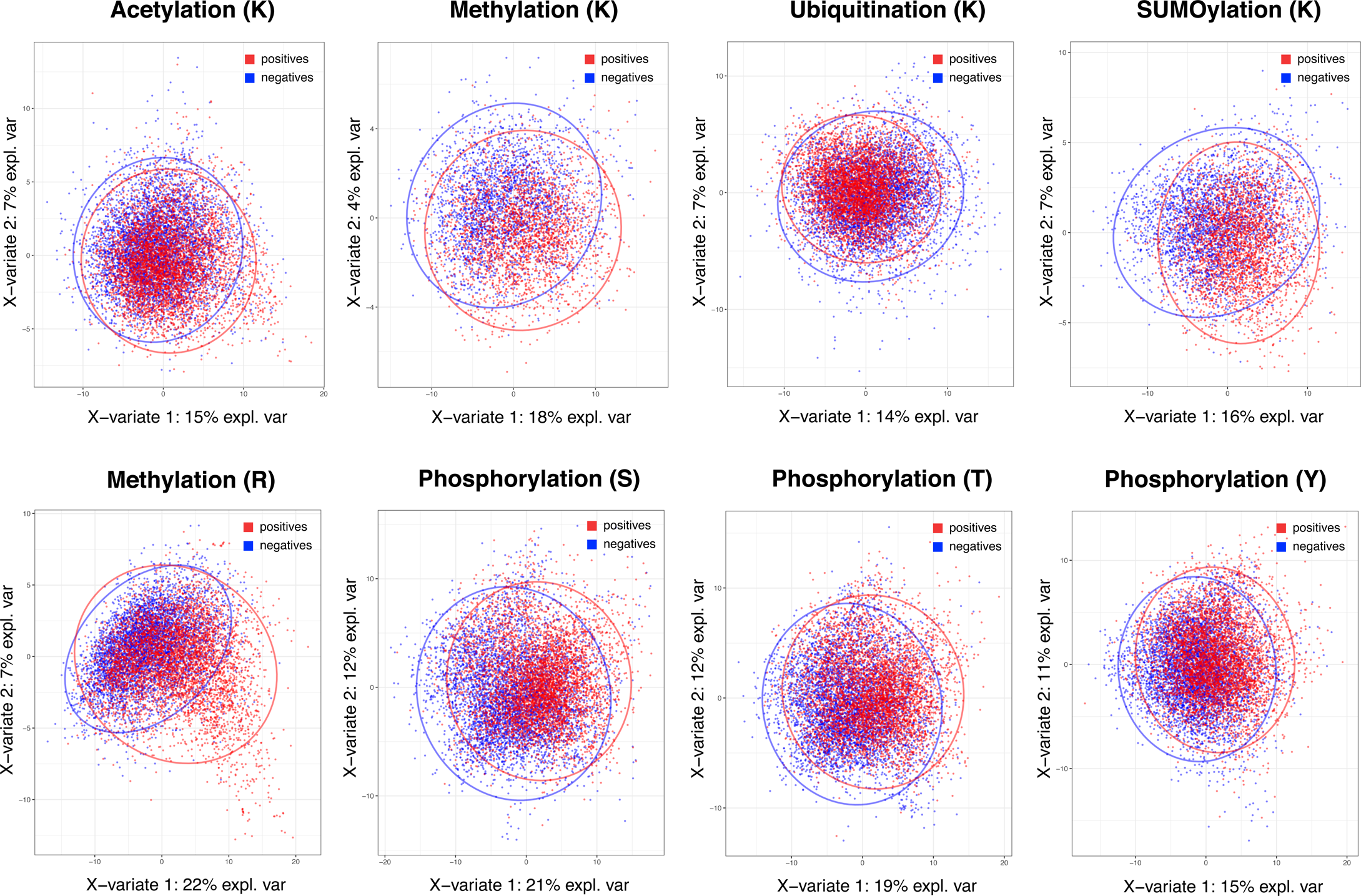
**A.** Sequence logo plots of the motifs obtained from PSP reported sites and additionally predicted sites using ggseqlogo^51^ across the five different modifications. **B-F**. Amino acid index, accessible surface area, half-sphere exposure, probability of coil and helix, and PSAAP information obtained from known and additionally predicted sites. The hierarchical clustering of five modifications was performed using all variables.

Several lines of evidence supported the validity of the PTMscape predictions. First, sequence motifs calculated from PTMscape predicted sites were similar to those calculated from experimental data alone (**Supplementary Figure 2**). Further, the protein domains showed a similar enrichment in modifications arising either from PTMscape predictions or experimental (PSP) data, with the exception of small differences for domains enriched for lysine/arginine methylation and lysine SUMOylation (**Supplementary Figure 3**). This difference is perhaps due to the low experimental coverage of these modifications. Therefore, we are confident that PTMscape provides accurate and biologically relevant predictions for protein modification sites to complement available experimental data.

Next, we examined modification-enriched proteins and their domains in detail. To do so, we mapped experimental and newly predicted sites to 487 most frequent protein domain families as defined by the Pfam database^26^. More than 159 of these domain families showed statistically significant enrichment of specific types of modifications (*q*-value < 0.05). **Supplementary Figure 4A** shows the heatmap of domain enrichment scores of all five modifications (-log10 of *p*-values). The domain families with the most statistically significant enrichment include protein kinase domain, zinc finger-C2H2 domain, RNA recognition motif, and histone domain (**Supplementary Table 3**). Protein kinases are often phosphorylated themselves as part of a signal transduction cascade, however, other modifications, such as ubiquitination, as also well-known to affect the kinase’s function.^27^ Both zinc finger domains and RNA recognition motifs bind nucleic acids, and abundant SUMOylation and phosphorylation events have been described.^28^’ ^29^ Finally, the histone modification ‘code’ is well-known, covering methylation, ubiquitination, and acetylation, for example. ^30, 31^

We then tested the enrichment of biological functions in the proteins harboring in-domain modifications (GeneMania, *q*-value < 10^-20^ only).^32^ **Figure 3A** shows that mRNA splicing, spliceosome, RNA processing were the most significantly enriched in the domains with modifications of lysine and arginine residues, i.e. SUMOylation, ubiquitination, and methylation – consistent with the enrichment in nucleic acid binding domains. In comparison, functions in signal transduction, DNA damage response, regulation of cell cycle, immune response, and growth factor responses were enriched in domains with a variety of modifications and affected different residues. Proteins whose domains accumulated threonine and tyrosine phosphorylation sites, and also lysine ubiquitination and acetylation often function in regulation of cell cycle, various receptor signaling pathways, and protein kinase activities – consistent with the abundant role of phosphorylation in signaling cascades. This landscape illustrates the co-existence of different modifications on the same protein, such as those for p53 mentioned above, and these proteins can have many essential biological functions. We further explored the modifications that co-occurred in *local neighborhoods* of protein sequences in a later section.

**Figure 3.**
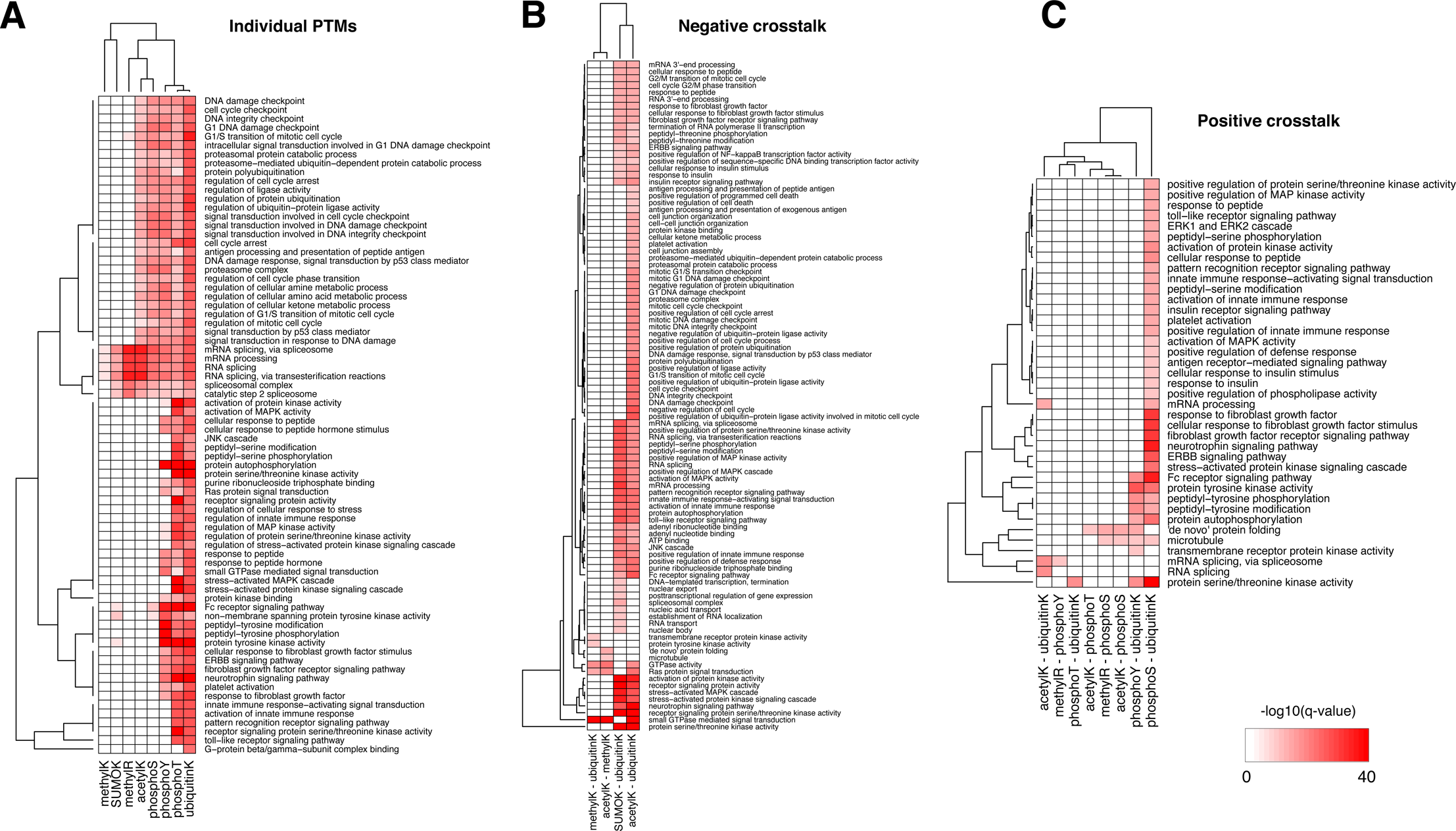
Biological functions enriched in (**A**) individual modification sites, (**B**) negative crosstalk, and (**C**) positive crosstalk events. The color of heatmap was -log10 of *q*-values reported from GeneMania.

### Highly predictive features of individual modifications

In addition to its robust performance, the advantage of PTMscape’s linear SVM is the straightforward inference of the most predictive properties for specific modification types, as extracted from the 173 total predictors. This interpretation is possible because the linear kernel in SVM requires that the decision boundaries be linearly associated with each variable, therefore naturally allowing us to describe the variables as positively or negatively associated with the modification.

**Figure 2 (B-E)** shows weight coefficients of the SVM predictors for all modification types. The amino acid indexes were the strongest predictors for phosphorylation, ubiquitination, acetylation and methylation (**Figure 2B**). In general, high hydrophobicity, partition coefficients, low solubility, low probability to be in an alpha helix/helix terminus, and propensity to be in a domain linker tended to be positively predictive of these modifications. These descriptors are often found in membrane proteins and link to a biased amino acid composition. However, other predictors varied for the different modifications. For example, hydrophobicity and amino acid composition of intracellular proteins were positively predictive only for phosphorylation, but in no other modifications. By contrast, the propensity of amino acids to be positioned in beta sheets and the properties associated with helical ends (e.g. chain reversal) were positively predictive for several lysine modifications, such as SUMOylation, ubiquitination, and acetylation.

Solvent accessibility and secondary structure showed different patterns for different modification types (**Figure 2C**). For example, tyrosine – but not serine or threonine - phosphorylation was more likely in sequences with poor solvent accessibility, i.e. in a pocket shape. This result is consistent with the known higher similarity of serine and threonine phosphorylation compared to tyrosine phosphorylation.^33^ Lysines modified by ubiquitin, SUMO or acetylation were negatively associated with solvent accessibility. Further, the residues in coiled coils tended to be phosphorylated, arginine methylated, or SUMOylated (**Figures 2D and 2E**). Residues in helices tended to be acetylated, methylated, or SUMOylated.

Lastly, PTMscape identified sequence motifs matching those known from literature. The corresponding PSAAP scores were predictive for those modifications with clear sequence motifs, i.e. lysine and arginine methylation, SUMOylation, and serine/threonine phosphorylation. The sequence logo plots for the experimentally detected and newly predicted sites show the relative strength of the motifs (**Figure 2A**). The best examples are the ΨKxE consensus motif for SUMOylation^34^ and GAR consensus motif for arginine methylation,^35^ and proline at +1 position of serine/threonine phosphorylation.^36^ By contrast, there were no clear consensus motifs that could *globally* predict ubiquitination, lysine acetylation, and tyrosine phosphorylation, as has been demonstrated in previous work.^33, 37^

### A global map of candidates for modification crosstalk

PTMscape’s unified, proteome-scale prediction scheme allows us to move towards systematic analysis of modification hotspots and their possible causal interactions. Such interactions among modifications are also known as crosstalk.^38^ Here, we define positive crosstalk as instances in which several modifications of the same or different type localize to the same sequence region (within five amino acids), but they do not affect the same residue. Many of these modifications might occur simultaneously, e.g. ubiquitination often affects multiple lysine residues within the same structural neighborhood. Positively ‘interacting’ modifications might also have temporal or even causal relationships: one modification might trigger another modification of the same protein within the same sequence neighborhood. For example, phosphorylation often triggers subsequent SUMOylation, e.g. as is found for heat shock transcription factor HSF1for its serine and lysine residues at positions 303 and 298, respectively.^39^ The p53 tumor suppressor protein can be activated through phosphorylation of serine 46, which promotes acetylation at lysine 382 through positive crosstalk. ^40, 41^

In comparison, we define negative crosstalk as events in which *the same residue* is targeted by multiple modifications. In our study, such negative crosstalk can occur only for lysine residues: for example, lysine can be acetylated, methylated, SUMOylated, or ubiquitinated. However, since these modifications are chemically largely exclusive, we assume that only one can occur at a given time. Therefore, negative crosstalk represents cases in which the modifications ‘compete’ for the same residue in a temporal or causal manner. Such negative cross occurs, for example, for lysine 382 in p53 mentioned above in which acetylation then competes with methylation.^42^

Using the combined set of experimental and predicted modifications derived from PTMscape and the PSP database, we first examined hotspots of potential negative crosstalk. For example, there are 532,420 lysine residues in total across the human proteins considered here, which are candidates for ubiquitination, SUMOylation, acetylation, and methylation. Of these, 9,511 lysines (1.8%) had a PTMscape score above the threshold for two or more different modifications, i.e. their sequence context suggested that indeed, multiple modifications compete for the same lysine. As neither PSP nor PTMscape provides temporal resolution for the different modifications, it remains to future research to resolve their causal relationships.

We then tested for enrichment of these negative crosstalk sites with respect to common protein domains and biological functions (**Supplementary Figure 4B, Supplementary Table 4**). Histone and tubulin domains showed the most pronounced enrichment of lysine residues with multiple modifications. Further, several zinc finger domain families and the RNA recognition motif were enriched in negative crosstalk sites for SUMOylation and ubiquitination. Proteins harboring negative crosstalk for acetylation and ubiquitination were involved in processes including DNA damage response, RNA processing, immune response, and signaling cascades (GeneMania q-value <10^-10^); the proteins with negative crosstalk for SUMOylation and ubiquitination specifically had enrichment of mRNA processing and kinase signaling pathways (**Figure 3B**).

Positive crosstalk, i.e. the statistically significant accumulation of several modifications within the same immediate sequence neighborhood, was much more frequent than negative crosstalk. Since testing all possible combinations of modifications is statistically infeasible, we focused on evolutionarily conserved pairs of modifications^43^. **Supplementary Figure 4C** shows the landscape of complex interplay of 80 domain families with statistically significant enrichment of their combined occurrence (Fisher exact test, *q*-value < 0.05, see **Methods**). When testing for enrichments for combinations of phosphorylation, acetylation, and ubiquitination, we found that positive crosstalk events had similar biases as the individual modifications, i.e. they were enriched in domain families such as histone and linker histone, ubiquitin, tubulin, protein kinase domains, and the RNA recognition motif, and in functions such as mRNA processing and RNA splicing (**Figure 3C**, GeneMania *q*-value < 10^-10^). Protein kinase activity, regulation of peptide hormones, receptor signaling pathways were enriched in the crosstalk sites among phosphorylation, acetylation, ubiquitination and SUMOylation events. The full landscape of domain-level and function-level enrichment of crosstalk is provided in **Supplementary Table 4**.

## Discussion

In this work, we developed a new computational tool called PTMscape for generic, unified prediction of protein modifications at whole-proteome scale. To the best our knowledge, PTMscape is equipped with the most comprehensive set of predictors and uses a fast and robust machine-learning algorithm whose results are easily interpretable for their biological meaning. We demonstrate PTMscape through the example of five post-translational modifications analyzed across the *entire human proteome.*

PTMscape moves beyond existing tools for *in silico* prediction of modifications in several ways. First and most importantly, it is generic and can be used for any type of the ~200 currently known modifications as long as there is training data, such as the data from a single mass spectrometry analysis with enrichment of the given PTM. Second, PTMscape is easy to use at proteome scale as it is installed locally. This feature also allows the user to customize the analysis with respect to the types of modifications that are analyzed, the background sequence file, the types of predictors to be evaluated and, most importantly, the experimental data that is used for training. Therefore, PTMscape provides a tool for the expert user who needs a large-scale, comprehensive analysis in the most flexible format. Finally, PTMscape provides substantial additional information that will help interpretation of the results. It evaluates the modification’s microenvironment, including the physicochemical properties, site-specific secondary structure properties, and motifs within the sequence neighborhood. PTMscape also includes less commonly used predictors, such as the accessible surface area or secondary structures. Including analysis of such features of the protein three-dimensional structure proved to be highly valuable in particular for modifications such as ubiquitin that lack a defined sequence motif.^44^’ ^45^

These aspects also illustrate future extensions that address some of PTMscape’s current limitations. For example, for statistically robust analysis, a few thousands of experimentally detected sites need to be available. In particular, for modifications that target a wide range of sequences without the presence of a global motif, such as ubiquitin mentioned above, the training data needs to include a wide range of diverse sites. However, for many modifications, such as O-linked glycosylation, available data is limited at the moment. Therefore, future extensions of PTMscape will address this challenge by exploring additional predictive feature variables extracted from external data, such as information on protein-protein interaction data or features pertaining to each residue’s position in the protein structure as obtained from experiments or modeling – moving beyond the current sequence-based prediction. Similarly, future extension might also use windows surrounding possible modification sites based on structural similarity, rather than sequence proximity, or pursue simultaneous prediction of such multiple modifications in a single model, incorporating the domain and other structural information.^6, 46^

## Methods

### Overview of the workflow

PTMscape is a command line-based tool written in R (http:/cran.r-project.org), and it is provided through a GitHub repository, at https://github.com/ginnyintifa/PTMscape under Apache 2.0 license. PTMscape builds sub-sequence windows centered at known or candidate PTM sites. It encompasses a comprehensive set of features describing modification events, which are used as training and test data in the linear kernel SVM library called *liblinear*^47^. The prediction output is further annotated with domain and sub-cellular information. Finally, PTMscape offers statistical tests for the enrichment of protein domains in individual PTM sites and co-occurrence of any two types of PTM events (crosstalk).

### Source data

From Uniprot Swiss-Prot database, we downloaded 20,201 canonical human proteins (as of August 2017).^5^ Experimentally validated PTM sites corresponding to five PTM types (**Table 3**) were gathered from PhosphoSitePlus database^3^. A total of 538 numerical indices representing various physicochemical and biomedical properties for 20 amino acids were retrieved from AAindex.^18^ Position specific secondary structural features, accessible surface area (ASA) and half sphere exposure (HSE) of each residue were predicted with the SPIDER3 tool.^19^ Information on protein domain families were obtained from the Pfam using the command line tool.^26^

### Removal of sequence redundancy

To obtain a set of non-redundant proteins for classifier performance evaluation, we used the CD-HIT software^48^ at similarity level 0.3 to reduce the redundancy of the whole proteome (sets of proteins containing at least one modifiable residue for respective modification). As a result, prediction of phosphorylation on serine (S) site was conducted for 11,490 non-redundant sequences which contained at least one serine. Similarly, we have 12,188, 11,986, 12,222 and 11,872 nonredundant sequences for analysis of PTMs on threonine (T), tyrosine (Y), lysine (K), and arginine (R) respectively.

### Amino acid index dimension reduction

To reduce feature dimension and minimize correlation within the indices, we hierarchically clustered the features downloaded from amino acid index into 53 clusters using dynamic tree cut algorithm^49^, each summarizing a group of similar physicochemical properties for the amino acids such as hydrophobicity, eccentricity size, solubility etc. Detailed information on the clusters is described in **Supplementary Table 5**. For each window, we calculated the average property of each cluster (excluding the center site). Note that all other features below were position specific around the modified residue (-12, -11, …, -2, -1, +1, +2, …, 12).

### Position specific structural features

SPIDER3 is a sequence-based prediction tool of local and non-local structural features for proteins using Long Short-Term Memory Spider.^19^ Prediction output includes probabilities of three secondary structures (helix, strand, and coil), ASA and HSE. For each flanking residue in a window, we extracted four features, namely p_Coil, p_Helix, ASA and HSE. Under the default mode, we constructed 96 dimensional position-specific structural features.

### Position specific amino acid propensity (PSAAP)

PSAAPs were derived from the frequency of each amino acid in each of the 24 positions within all the windows centered with positive PTM sites (excluding the center residue). Therefore, the matrix is composed of 20 rows and 24 columns. Each column records the probability of 20 amino acids appearing in the corresponding position based on the observed frequency in the positive sites. For instance, for position *j*, column *j* can be expressed in the following vector:

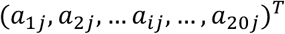

a_*ij*_ represents the probability of amino acid *i* appears in the *j*-th position.

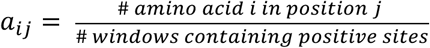

Note that 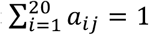, *for j* = 1, 2, … 24.

### Feature data generation

The data generation module maps protein identifiers and positions of experimentally determined PTM sites (positive sites) to any protein sequences in the user-provided FASTA file, and computes average AA specific features for the *k*-mer window centered surrounding each candidate site (default *k* = 25, having 12 amino acids on both sides of each site). The feature variables used in the default mode include 53 clusters of physiochemical properties derived from the AAindex database, 96 (24x4) position specific structure properties generated by SPIDER3, and 24 dimensional features formed by position specific amino acid propensity (PSAAP). We refer to the resulting data as the feature data. The program also generates feature data for windows that contain negative sites, i.e. sites without experimental evidence for their modification.

### Support vector machine with linear kernel

PTMscape uses support vector machine (SVM)^50^ with linear kernel, to achieve the best classification of modification sites while avoiding over-training. Moreover, linear SVM provides easy interpretation of how each property contributes to the decision function of the SVM. In a linear kernel setting, the classification is achieved by solving the following problem:

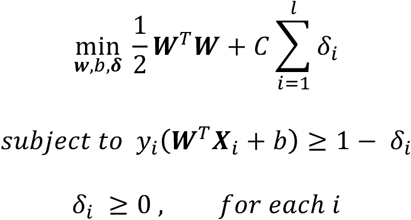

where *y_i_* equals to 1 or -1 representing the class label of support vector **X**_*i*_, ***W*** represents the weight coefficient for each feature. *C* is the soft margin tuning parameter. The model fitting is performed using *liblinear* library. During training, we used L2-regularized L2-loss support vector classification (-s 2). To obtain the probability score in addition to the classification labels from each prediction, we modified the code of *liblinear* as suggested in the FAQ page (https://www.csie.ntu.edu.tw/~cjlin/liblinear/FAQ.html).

### Prediction performance metrics

At a given score threshold, we recorded the true positives (TF), true negatives (TN), false positives (FP) and false negatives (FN). The performance metrics were defined as:

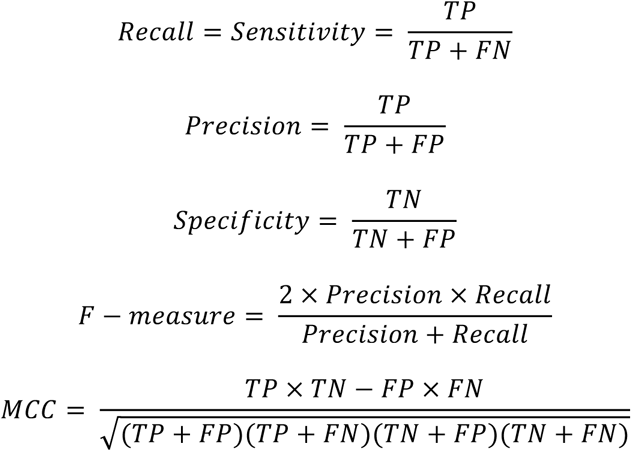

where MCC denotes Matthew’s correlation coefficient.

### Evaluation of the classifier performance with 10-fold cross validation

To evaluate the performance of the classifier unbiasedly, we implemented a 10-fold cross validation scheme. For each PTM type, we first randomly sampled the same number of negative and the positive windows and randomly divided both sets (positive and negative) into ten folds. Each time we assembled nine out of the ten folds as a training dataset and used the remaining one fold as a validation dataset. Therefore, training and testing was performed on independent datasets. The performance metrics such as Area Under Curve (AUC) and Matthew’s Correlation Coefficient (MCC) were calculated by taking the average of the ten iterations.

### Predictions with PhosphoSVM and ESA-Ubisite

To benchmark our tool against PhosphoSVM, we used the data retrieved from their web portal at http://sysbio.unl.edu/PhosphoSVM/download.php. PTMscape extracted features for 9,688, 2,919, and 1,269 positive PTM sites (on 2,545, 1,499, and 805 proteins) for serine, threonine and tyrosine phosphorylation, respectively. Then 10-fold cross validation was conducted on the feature data generated. For comparison with ESA-UbiSite, we obtained the data consisting of 645 positive sites and 10,336 non-validated sites from 379 protein sequences from their work^17^. Specifically, we collected the proteins in the UbiD set for which position specific structural features are available. PTMscape was applied to 355 proteins with 613 positive ubiquitination sites and the same proportion of training and test division was implemented. For model training, we obtained the same number of negative windows as positive windows by randomly sampling the negatives from the whole negative set.

### Comparison with ModPred

ModPred predicts as many as 23 different modification types in different species. The predictor in ModPred is trained with sequence-based properties, physicochemical features and evolutionary conservation information, where ensembles of logistic regression models were built per modification residue type. Using the stand-alone version ModPred_Linux64, we predicted all PTMs on lysine residues for the proteins in the Swiss-Prot database that are longer than 30 residues and hold one or more modifiable lysine (without PSSM features). The AUC of ModPred predictions was calculated by comparing the output prediction scores with known PTM status from the PhosphoSitePlus database.

### Prediction on the whole human proteome

To predict PTM sites in the whole proteome, we gathered all 17,612 sequences for which position-specific structural features can be extracted by SPIDER3. PTMscape was applied in a two-fold cross-prediction scheme, where we used half the data as training data and predicted on the other half, and switched the role of training and test data to make predictions in the former. This ensures that the same data point is not used for training and prediction simultaneously, rendering predictions unbiased. During the training, we randomly sampled from the negative set in a way that the sizes of positive windows and negative windows were the same.

All predicted sites are provided through the website http://137.132.97.109:59739/CSSB_LAB/. The tables list the modification type, PTMscape score, the modification type, the modification’s threshold for 99% specificity, and reports on significant enrichment in protein domain families or protein functions. The table also indicates whether a modification site is ‘new’, i.e. if it has not been observed in the PSP database that we used for training.

### Enrichment analysis of PTM occurrence in protein domains and biological functions

For the enrichment analysis of individual PTMs in each domain, we first counted the number of positive and negative PTM sites in the domain and across the proteome, and performed a chi-square test to test whether the frequency of a PTM type in a domain is significantly higher than the expected frequency across the proteome. We adjusted for multiple testing by selecting domain families with *q*-value smaller than 0.05.

For the analysis of negative crosstalk, we tested association of two competing modifications as follows. In each domain, we constructed a 2-by-2 contingency table where rows and columns are positive and negative status of the two competing PTM events on all modifiable sites within that domain family. We then tested whether the two PTMs are significantly more frequent in the domain than expected under by Fisher exact test.

For the analysis of positive crosstalk in a domain, we similarly constructed a contingency table, where the sum of the numbers in the four cells is the number of all pairs of modifiable residues within a domain. Here a modifiable pair of residues in a domain refers to two amino acids located within 5 amino acids, where one residue is modifiable in one of the two PTM types and the other residue is modifiable in the other PTM type. Each pair contributes to one of the four cells in the contingency table based on the PTM status of respective types. After the construction of the contingency table, we tested the significance of enrichment of co-occurring two PTMs in a domain by fisher exact test. These significance scores (*p*-value) were further adjusted for multiple testing (*q*-value), and the domains with *q*-value smaller than 0.05 were considered to have a significant positive crosstalk event.

Significant crosstalk events are listed in **Supplementary Tables 2 and 3**. The tables list the paired modifications, the p-values from the chi-squared test and the q-values as multiple testing adjusted significance measure.

### Function enrichment analysis of individual PTMs and crosstalk

A list of proteins with individual and crosstalk events in protein domains was analyzed with Genemania^32^ for function enrichment. In GeneMania, we used 2017-07-13-core version of the human data, with annotation limited to Pathway only, without finding any other top related genes.

## Acknowledgments

This work was supported in part by a grant from the Singapore Ministry of Education (to HC; MOE2016 T2-1-001) and the United States National Institutes of Health (to CV and HC; 5R01GM94231).

## Availability

The software is freely available at https://github.com/ginnyintifa/PTMscape (Apache license), along with a tutorial and example data sets. Additional feature information for proteome-scale predictions is provided through a website at http://137.132.97.109:59739/CSSB_LAB/.

